# A phylogenomic framework for avian feather lice (Phthiraptera: Ischnocera)

**DOI:** 10.64898/2026.01.09.698582

**Authors:** Kevin P. Johnson, Julie M. Allen, Avery Szewczak, Kimberly K. O. Walden, Jorge Doña

## Abstract

Comprehensive phylogenies serve as foundations for taxonomic, comparative, and coevolutionary studies. Avian feather lice comprise the most diverse Parvorder (Ischnocera) of parasitic lice (Phthiraptera). Convergence and reduction of morphological features in this group have made past attempts at understanding the evolutionary relationships of feather lice challenging. Several recent phylogenomic studies have begun shedding light on the broader scale evolutionary tree of this group. However, additional taxonomic sampling is needed for a more complete understanding of the phylogeny and for higher statistical power in comparative studies. Here we investigate the higher-level relationships of feather lice based on the most comprehensive taxon sample to date. We leverage genome sequences of 260 samples of feather lice and 25 outgroup taxa to reconstruct a phylogenomic tree based on 2,395 target nuclear ortholog genes, using both concatenated and coalescent methods. These trees provide high support across nearly all branches of the topology, resolving backbone relationships as well as more detailed relationships within major groups. These phylogenies provide a framework for future comparative studies of feather lice, including comparisons with avian phylogeny and investigating patterns of diversification.

## Introduction

Avian feather lice (Phthiraptera: Ischnocera) have become a model system for understanding the interplay between ecological interactions and coevolutionary diversification with their hosts (Clayton et al. 2015). Over 3,000 species of feather lice have been described (Price et al. 2003), and many more are likely to be discovered (Gustafsson and Bush 2017), making this group an important component of insect diversity. These ectoparasites spend their entire lifecycle on the body of their avian hosts (Clayton et al. 2015), which in part contributes to their generally high degree of host specificity (Price et al. 2003). Members of Ischnocera consume the feathers of their hosts, which can interfere with thermal regulation and can impact host survival (Clayton et al. 1999) and reproductive success (Clayton 1990). In response, birds combat these ectoparasites by preening and scratching, which are effective in removing lice (Clayton et al. 2005). Feather lice have diversified into several ecomorphs, which differ in the way they escape these host defenses (Clay 1949, Johnson et al. 2012, Kolencik et al. 2024). These interactions set up coevolutionary dynamics that make feather lice an excellent system to integrate both experimental and comparative approaches.

Comparative studies require well supported phylogenetic trees with a high level of taxon sampling at the scale at which the comparisons are being made. Generic diversity of feather lice generally corresponds to ecomorphology and host association with avian orders or families (Price et al. 2003). Thus, uncovering broad patterns of codiversification requires sampling at these scales. In the past, however, understanding of the evolutionary relationships among feather lice was hampered by their extreme morphological diversification and convergence into ecomorphs (Clay 1949, Johnson et al. 2012). This morphological complexity above the level of genus hindered higher level classification of this group, and even generic level keys were not forthcoming (Smith 2001, Gustafsson et al. 2025). Phylogenetic studies based on morphology alone tended to group genera of the same ecomorph together (Smith 2001). However, early molecular data sets from Sanger sequencing revealed that these groupings were largely driven by morphological convergence across ecomorphs (Johnson et al. 2012). Nonetheless, given the high rate of molecular evolution of lice, molecular phylogenetic studies of just a few genes were not able to resolve with confidence the higher level relationships of feather lice (Cruickshank et al. 2001, Johnson and Whiting 2002, Johnson et al. 2003, Johnson et al. 2012).

Recent advances in phylogenomic approaches have made possible new understanding of higher level evolutionary relationships of both these parasitic insects (Johnson et al. 2018, 2022, de Moya et al. 2019, de Moya 2022) and their avian hosts (Jarvis et al. 2014, Prum et al. 2015, Feng et al. 2020, Stiller et al. 2024). These studies have leveraged whole genome sequencing to achieve data sets of hundreds or thousands of genes for phylogenetic analysis. Phylogenomic trees have revealed several major clades of feather lice, with high support and consistency for the relationships among these clades. Although there is still some uncertainty in relationships among enigmatic genera near the base of the tree (Johnson et al. 2018, de Moya et al. 2019, de Moya 2022), these studies have provided new insights into the relationships among avian feather lice.

The goal of the current study was to develop the largest phylogenomic data set of feather lice (Ischnocera) to date, more than doubling the number of loci examined and nearly doubling the number of included terminals of feather lice over prior studies. This taxon sample not only includes representatives of as many genera as possible, but also includes a diversity of host associations corresponding to terminals in a major phylogenomic study of birds (Prum et al. 2015), which sampled 198 species of birds mainly focused on representing avian families.

Together these phylogenies will facilitate future direct cophylogenetic comparisons between birds and feather lice and enable comprehensive comparative studies at this taxonomic scale. To achieve this, we sequenced whole genomes from 260 representatives of feather lice comprising 131 genera. From these genomic read data sets, we assembled a target set of 2,395 single copy nuclear protein coding orthologs into a phylogenomic dataset. This dataset was analyzed using both concatenated and coalescent phylogenomic approaches to estimate the phylogenetic tree of feather lice. We also discuss and explore the implications of this feather louse tree as it relates to patterns of host association and biogeography.

## Methods

### Taxon Sampling

As an ingroup, we sampled 260 individual feather lice (Ischnocera) representing 131 genera (over 85% of described genera, as defined in Price et al. 2003, Gustafsson and Bush 2017). Our sample was designed to represent as many feather louse genera as possible and also to be matched to the avian families sampled by the phylogenomic study of Prum et al. (2015) to the extent possible. This sample comprises the most comprehensive sample of feather lice in a phylogenetic study to date. Outgroup samples comprised representatives of the Parvorders Amblycera (10 species), Trichodectera (4 species), Rhynchophthirina (1 species), and Anoplura (10 species). Samples were photographed as vouchers and deposited in Figshare (10.6084/m9.figshare.29266997). We identified each specimen to genus based on the illustrations and keys in Price et al. (2003). Some samples had been included as part of prior studies and we applied the determinations on identification made in these prior studies (Johnson et al. 2018, 2021, Johnson and Doña 2024, de Moya et al. 2019, de Moya 2022, Doña and Johnson 2023, Virrueta Herrera et al. 2020).

### Sample Preparation and Genome Sequencing

Specimens were stored in 95% ethanol at -80 C. Individual lice were selected for sequencing and washed in a vial with clean 95% ethanol. A Qiagen QIAamp DNA Micro Kit was used for total genomic DNA extraction from individual lice, which were first ground with a plastic pestel in a 1.5 ml tube in extraction buffer. Extraction followed manufacturer’s protocols, except that the initial digestion with proteinase K was performed for ∼24 to 48 hours. Extractions were eluted in 50 ul AE buffer and quantified using the high sensitivity kit with a Qubit 2.0 Flourometer.

From these total gDNA extractions, libraries were prepared with the Hyper library construction kit (Kapa Biosystems). These libraries were sequenced to produce 150bp paired-end reads of ∼400bp insert multiplexed to 48 libraries per S4 lane of Illumina NovaSeq 6000. All libraries were tagged with unique dual-end adaptors. This level of multiplexing was chosen with the aim of achieving at least 30-60X coverage of the louse genome assuming a genome size of approximately 200Mbp (Baldwin Brown et al. 2021, Sweet et al. 2023). Fastq files were generated by adaptor trimming and demultiplexing using bcl2fastq v2.20. We deposited the raw reads for each library in the NCBI SRA database (Supplemental Table 1).

### Gene Assembly

For compatibility with prior phylogenomic studies of lice and other hemipteroid insects (Johnson et al. 2021, 2022, Johnson and Doña 2024, Doña and Johnson 2023), we selected a target set of 2,395 ortholog genes. These genes were assembled from raw reads using aTRAM 2.0 (Allen et al. 2018). Prior to aTRAM gene assembly, *fastp* v0.20.1 (Chen et al. 2018) was used for further adaptor and quality trimming (phred quality 30 or more). For aTRAM assembly (iterations = 3, max-target-seqs = 3000), we used *tblastn* searches of the target genes and assembled the hits with the ABySS assembler. The Exonerate (Slater and Birney 2005) pipeline in aTRAM was used to stitch the exons sequences for each gene.

### Phylogenomics

Alignment of orthologs according to inferred amino acid sequences was performed using MAFFT (Katoh et al. 2002, Katoh and Standley 2013). We trimmed each aligned gene sequence using trimAL v1.4.rev22 (Capella-Gutierrez et a. 2009) using a 40% threshold for missing data. Alignments for each gene were analyzed using maximum likelihood (ML) in IQTree 2 v2.1.2 (Nguyen et al. 2015, Minh et al. 2020) using the inferred optimal model for each gene to obtain gene trees. Trees were rooted on Amblycera based on results of prior studies (Johnson et al. 2018, de Moya et al. 2019). Gene alignments were also concatenated into a supermatrix and analyzed in IQTree using a partitioned analyses. The optimal model for each partition was used in a ML search of the concatenated data set. Ultrafast bootstrapping with UFBoot2 (Minh et al. 2013, Hoang et al. 2017) was used to assess branch support. A coalescent approach to estimating the species tree was performed on the individual gene trees using ASTRAL-III (Zhang et al. 2018). Local posterior probabilities for each node in the coalescent tree were also calculated using this software.

### Biogeographic Reconstruction

To obtain an ultrametric tree for biogeographic analyses, we performed a dating analysis using the least squares methods (LSD2) in IQ-TREE 2 (To et al. 2016). We used the same calibration points from previous molecular dating studies of lice, including the divergence between human and chimpanzee lice (5–7 Ma), the divergence between Old World primate lice and those of Great Apes (20–25 Ma), and a fossil-based minimum age of 44 Ma for Menoponidae. Also, following prior studies, we rooted the tree at 92 Ma, corresponding to the split between Amblycera and other parasitic lice (Johnson et al. 2018, 2022).

Because only single samples of lice were used, we coded each louse according to the biogeographic distribution of its host species in its native range. Broad geographic regions of New World, Eurasia, Africa, Australasia, and Antarctica were used to encompass broad patterns of avian (and by association feather louse) biogeography. Using this coding and the ultrametric louse tree from the molecular dating analysis (above), we employed the R package BioGeoBears v1.1.2 (Matzke 2013, 2014) to estimate ancestral biogeographic distributions for each node in the feather louse tree. Outgroups were not included in this analysis because they were less densely sampled than the ingroup. We ran the DEC, DIVALIKE, and BAYAREALIKE models in BioGeoBears, both with and without the J (jump) parameter and selected the model with the lowest AIC score to estimate the best maximum likelihood reconstruction.

## Results

### Genome Sequencing and Gene Assembly

We obtained between 37 and 240 million total 150bp paired-end reads (Read 1 + Read 2) from each louse sample. The results from aTRAM assemblies of the 2,395 target genes produced between 1,327 and 2,358 (mean = 2,323) genes for each taxon. After alignment and trimming, 2,377 genes were retained for phylogenomic analyses with a total of 3,820,401 aligned base pairs in the concatenated matrix.

### Phylogenetic Relationships

The partitioned concatenated (Figure 1) and coalescent trees (Supplemental Figure 1) were highly similar, with only 21 of 259 ingroup nodes differing between them. In the respective trees, most ingroup branches received 100% bootstrap support (251/259 partitioned concatenated) or 1.0 local posterior probability (250/259 coalescent). In both trees, Ischnocera was recovered as monophyletic with maximum support (100% bootstrap, 1.0 local posterior probability). Both trees revealed several major clades within feather lice, the relationships among which were highly supported and consistent between the two analyses. The only difference between the two regarding rearrangement of major clades involved the placement of the genus *Columbicola* near the base of the tree. In the concatenated tree, this genus was sister to all other Ischnocera, while in the coalescent tree, it was sister to the clade containing the genera *Austrogoniodes*, *Craspedonirmus*, and *Carduiceps*. Otherwise, the broad backbone of the tree was completely stable between the two analyses.

**Figure 1.**
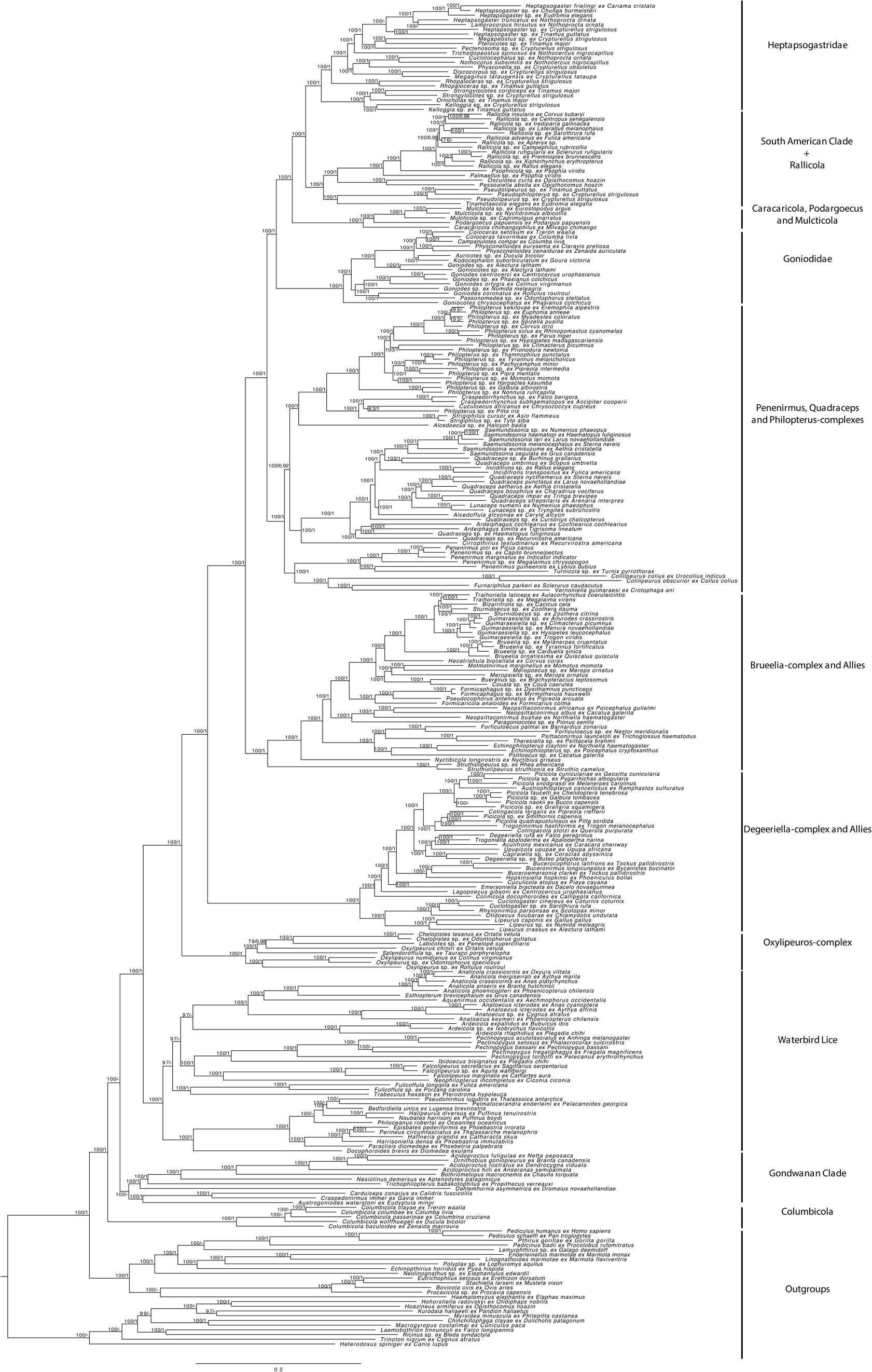
Phylogeny from partitioned concatenated analysis of 2395 target nuclear gene orthologues for ischnoceran feather lice and relatives. Numbers on branches indicate support from ultrafast bootstrap replicates and coalescent local posterior probability. Major clades are annotated on the right.

Notwithstanding the unstable placement of *Columbicola*, both trees agreed on an early diverging clade containing generally enigmatic genera with primarily Gondwanan distribution. The only genus of Ischnocera occurring on mammals (*Trichophilopterus* from Malagasy lemurs) is placed within this group. Generally, the genera in this clade are deeply divergent and are divided into two groups. The first contains the genera *Dahlemhornia* (from emus), *Trichophilopterus* (from lemurs), *Nesiotinus* (from large-bodied penguins), *Bothriometopus* (from screamers), *Acidoproctus* (from large-bodied waterfowl), and *Ornithobius* (from geese and swans). The second group in this clade includes *Austrogoniodes* (from penguins), *Carduiceps* (from shorebirds), and *Craspedonirmus* (from loons). The relationships within each of these two subclades are identical between the concatenated and coalescent trees.

The next major clade to diverge within Ischnocera is a group of lice occurring mainly on waterbirds, ranging from seabirds, rails, cormorants, herons, flamingos, grebes, and ducks. While several of these host groups have been shown to be related (forming a large clade of waterbirds) by phylogenomic studies of birds (Jarvis et al. 2014, Prum et al. 2015), others, such as ducks, are much more distantly related. Within this broader clade there are two subclades identified by both analyses. The first includes wing lice of tube-nosed seabirds (i.e. lice of the *Philoceanus*-complex Ledger 1980), but also the genus *Docophoroides*, a head louse of this host group. The second subclade includes a variety of genera that include wing lice (*Fulicoffula*, *Pectinopygus*, *Ardeicola*, *Aquanirmus*, *Esthiopterum*, and *Anaticola*) and head lice (*Trabeculus*, *Neophilopterus*, *Ibidoecus*, and *Anatoecus*) of a variety of waterbirds, but also includes *Falcolipeurus*, a wing louse, of raptors. The relationships among genera within this second subclade differ between concatenated and coalescent analyses and indeed some of these branches receive less than 100% bootstrap support. In particular, the position of *Ardeicola* switches with that of *Trabeculus* + *Fulicoffula* between the two trees, otherwise relationships among other taxa are generally stable.

Moving up the tree, the next major clade is the *Oxylipeurus*-complex, whose monophyly is strongly supported being united by a long stem branch. Members of this group occur mainly on landfowl (Galliformes), but one genus *Splendoroffula* occurs on turacos, a group of birds distantly related to Galliformes. The genus *Oxylipeurus*, as defined by Price et al. (2003) is paraphyletic, and both trees indicate that *Splendoroffula*, *Labicotes*, and *Chelopistes* are embedded within this genus.

The next diverging clade is also united by a long stem branch and includes members of *Lipeurus*, and the *Otidoecus* and *Degeeriella*-complexes (Ledger 1980). Within this group, *Lipeurus* is monophyletic and sister to the *Otidoecus*-complex, and together these are sister to the *Degeeriella*-complex. The concatenated and coalescent trees are generally consistent regarding the relationships within this broader clade, with only two minor branch rearrangements between them. The *Degeeriella*-complex contains a diversity of genera occurring across a wide range of host groups. Within this group, the genera *Picicola*, *Degeeriella*, and *Cotingacola* do not appear to be monophyletic.

The next major clade to diverge includes *Struthiolipeurus*, *Nyctibicola*, parrot lice, plus the highly diverse *Brueelia-*complex (Ledger 1980). Within this major clade, the concatenated and coalescent trees are identical in topology and all branches are supported by 100% bootstrap and 1.0 local posterior probability. The genus *Struthiolipeurus* (occurring on ostriches and rheas) is sister to all other members of this clade. Parrot lice are divided into two distinct clades that form a grade with respect to the *Brueelia*-complex and allies (Johnson and Doña 2024). The genera *Formicaricola*, *Pseudocophorus*, and *Formicaphagus* form a clade that is sister to the *Brueelia*-complex, which is now recognized to contain at least 38 genera (Gustafsson and Bush 2017).

Further up the tree, a major clade including three diverse genera (*Penenirmus*, *Quadraceps*, and *Philopterus*; as defined in Price et al. 2003) is highly supported by both concatenated and coalescent analyses. Each of these genera has related taxa that together form broader complexes. Two of these, *Quadraceps* and *Philopterus*, form the basis of large generic complexes that are highly supported by both analyses. However, the placement and monophyly of the genus *Penenirmus* (as defined by Price et al. 2003) is less clear. In both trees the representative of the genus *Turnicola*, is sister to a sample of *Penenirmus* from an African barbet (*Lybius*). However, this result conflicts with a broader study of the genus *Penenirmus* (Johnson et al. 2021), which found *Turnicola* to be outside of this genus. In addition, the genera *Vernoniella*, *Furnariphilus*, and *Colilipeurus* may be near this group, but the details of the placement of these taxa differs slightly between the concatenated and coalescent trees. Several of these genera (*Vernoniella*, *Colilipeurus*, and *Turnicola*) are on very long branches, such that issues related to long branch attraction may be underlying the instability of these taxa. In contrast, the concatenated and coalescent trees are completely congruent for the *Quadraceps*-complex (*Cummingsiella*-complex sensu Ledger 1980), and only one minor terminal branch rearrangement is observed in the *Philopterus*-complex. As found by a recent study based on Sanger sequencing (Kolencik et al. 2022), the genus *Debeauxoecus*, from Old World pittas, is far outside of *Philopterus* even though it had previously been placed within it (Price et al. 2003).

Beyond this, there are four additional major clades that together form a major group. Two of these (Goniodidae and Heptapsogastridae) have sometimes been recognized as distinct families (Hopkins and Clay 1952, Smith 2000) and were previously thought to be “basal Ischnocera”. However, these two clades group together with two other clades: 1) one which includes the widespread genus *Rallicola* as well as other genera primarily from taxa in South America and 2) the other including the genera *Caracaricola* (from caracaras), *Podargoecus* (from forgmouths), and *Mulcticola* (from nightjars). The clade Goniodidae occurs on landfowl (Galliformes) and doves (Columbiformes), while Heptapsogastridae occurs exclusively on tinamous in the Neotropics. Both the concatenated and coalescent trees agree on the arrangement of these groups with Goniodidae being sister to the remaining three. Within these clades, the two analyses are in nearly complete agreement except for two minor terminal rearrangements within *Rallicola* and within Heptapsogastridae. Within Goniodidae, the lice of doves form a clade, but are embedded within those from landfowl. Lice of tinamous that are not members of Heptapsogastridae, also form a clade with the *Rallicola*-complex, but it appears that the head louse genus *Pseudophilopterus* is embedded within the wing louse *Pseudolipeurus*, making the latter paraphyletic.

### Biogeographic Patterns

Generally, genera of feather lice are restricted to single families or orders of birds. As such, their biogeographic distribution is highly influenced by the distribution of their hosts. However, several major clades of lice that group taxa from more distantly related hosts occur within restricted biogeographic realms. For example, as mentioned above, there is a clade of taxa for which several genera occur exclusively in Gondwanan regions, such as *Nesiotinus* in Antarctica, *Trichophilopterus* in Madagascar, *Bothriometopus* in South America. There are also major clades of lice that mostly occur in single biogeographic regions, such as Heptasogastridae plus the *Rallicola*-complex whose members occur primarily in South America. A biogeographic ancestral states reconstruction using BioGeoBEARS indicated that BAYAREALIKE + J was the best model. Under this model, the New World was the ancestral area with high likelihood for many of the deep nodes in the tree (Figure 2). This was followed by independent “jump” dispersal diversification into other biogeographic regions mainly between 10 – 30 million years ago, a time in which the continents were well separated, consistent with the notion that this was dispersal driven speciation.

**Figure 2.**
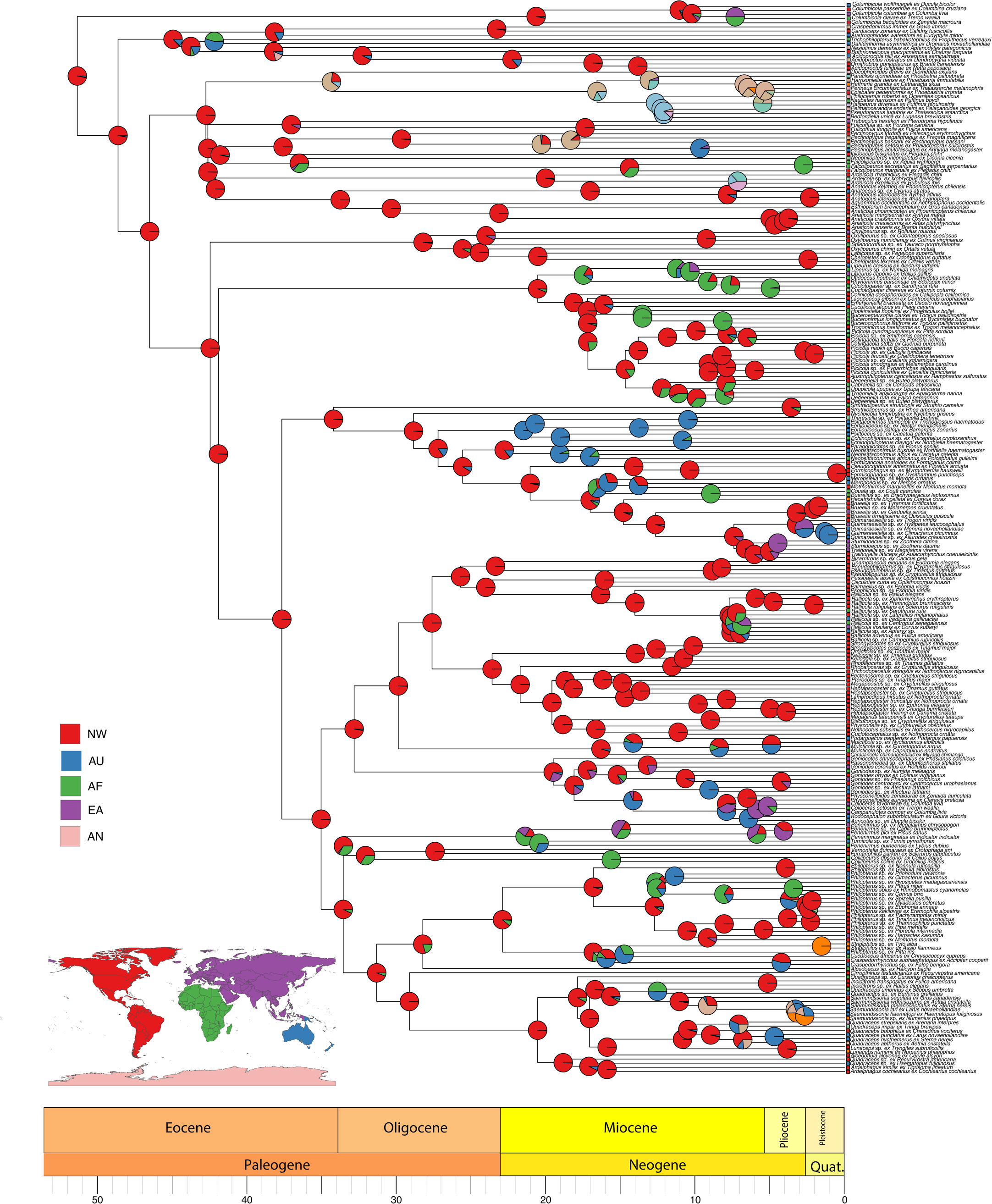
Dated phylogenetic tree of ischnoceran feather lice. Pie charts at each node indicate the likelihood that the ancestor occupied a particular biogeographic range, based on a historical biogeographic reconstruction with BioGeoBEARS under the BAYAREALIKE+J model. Colors correspond to major biogeographic regions shown in the map (NW = New World, AF = Africa, EA = Eurasia, AU = Australasia, AN = Antarctica). Squares at the tips indicate the current distribution of each taxon across biogeographic regions. For clarity, only the three most likely ancestral ranges are shown at each node, with their probabilities rescaled to sum to one. Additional mixed-area states are provided in Supplementary Figure S2.

## Discussion

Phylogenomic analyses of 2,395 target ortholog genes for 260 samples of feather lice (Ischnocera) and 25 outgroups revealed strongly supported and stable relationships throughout most of the tree. The vast majority of ingroup nodes were stable to method of analysis (only 21 of 259 nodes changed between concatenated and coalescent trees) and were supported with maximal support (251 of 259 ingroup nodes 100% bootstrap; 250 of 259 ingroup nodes 1.0 local posterior probability). These trees provide a new framework for broad-scale comparative and evolutionary studies of this major group of parasites.

The backbone structure of the tree was stable and revealed relationships among major well-supported clades of feather lice. These clades typically correspond to generic complexes of lice and relatives that are revealed by this phylogeny. The circumscription of these major clades has implications for taxonomy and could form the basis for new family level classification. Widespread convergent evolution in feather louse ecomorphology (Johnson et al. 2012, Kolencik et al. 2024) has limited the ability of prior studies to fully place many genera into a higher level framework. These limitations have led to the placement of all members of Ischnocera into a single family (Philopteridae) (Price et al. 2003). (Note, the mammal louse family Trichodectidae has now been shown to be outside of Ischnocera, instead being more closely related to other mammal lice [Johnson et al. 2022], a result also borne out by the current study). This classification of all feather lice into a single family stands in contrast to other major groups of lice, such as Anoplura, which is divided into 15 families (Durden and Musser 1994), even though it is much less diverse. Here we review the major groups of feather lice identified by the current study and the implications of these results for taxonomic and biological interpretations.

### Major Clades

#### Gondwanan Clade

Even though relationships at the very base of Ischnocera are somewhat unstable to method of analysis regarding the placement of the genus *Columbicola*, a group of additional enigmatic genera emerges together in a clade. Although most of the host lineages in this broader clade of lice are not closely related (Prum et al. 2015), many of these genera occur on birds with exclusively Gondwanan distribution, such as *Nesiotinus* and *Austrogoniodes* from penguins (Antarctic regions), *Dahlemhornia* from emus (Australia), *Bothriometopus* from screamers (South America), and the early diverging lineages of *Acidoproctus* from magpie geese (Australia), and whistling ducks (pan-Tropical). One somewhat unexpected result is that although the penguin lice *Nesiotinus* and *Austrogonides* both occur in this broader clade, they are not each other’s closest relatives, being somewhat distantly related. *Nesiotinus* occurs only on the very large bodied King Penguin (*Aptenodytes patagonicus*), while *Austrogoniodes* occurs broadly on nearly all penguins, including the King Penguin.

The one genus of Ischnocera that occurs on mammals, *Trichophilopterus*, is also in this clade, being found on Madagascan lemurs. Prior studies have indicated that the origin of *Trichophilopterus* was with high likelihood a host switch from birds to mammals (Johnson et al. 2022). Although *Acidoproctus* does occur on some whistling duck hosts that have some of their distribution in Madagascar, these lineages are relatively derived compared to *Trichophilopterus*. Rather, *Trichophilopterus* diverges near the base of this clade after the divergence of *Dahlemhornia* from emus. It is noteworthy that another ratite lineage now extinct, the elephant bird (*Aepyornis maximus*), occurred in Madagascar. Given that emus, another ratite from Australia, host lice of this lineage, it is conceivable that the elephant bird hosted a louse from this group and that this was the source of the host switch from birds to mammals in this case.

Although predominantly Gondwanan, there are also members of Arctic lineages in this group, particularly *Carduiceps* from shorebirds and *Craspedonirmus* from loons, which are sister taxa. Together these two genera are related to *Austrogoniodes*, which occurs on Antarctic and sub-Antarctic penguins. Although this geographic and host distribution might seem unexpected, the shorebird hosts of *Carduiceps* undergo very long-distance migrations between the Arctic and sub-Antarctic regions. This migratory behavior may have provided a means for lice to disperse from southern regions to the Arctic and also become established on loons, which breed in the Arctic. Similarly, the more derived lineages of *Acidoproctus* occur on migratory ducks and are sister to the genus *Ornithobius* from geese (all Northern Hemisphere), again indicating likely dispersal from Gondwanan to Northern Hemisphere regions by lice likely facilitated by host dispersal. The paraphyletic relationships of *Acidoproctus* with respect to *Ornithobius* indicate that generic reclassification is needed.

Although the exact placement of *Columbicola*, which occurs on doves throughout the world, is unclear, a detailed cophylogenetic study of this genus and their hosts (Boyd et al. 2022) revealed that doves also had origins in Gondwanan regions (particularly Australia and South America) and dispersal from these regions provided a means for *Columbicola* to switch to lineages in other regions of the world. The long stem branch of *Columbicola*, combined with the long branches of many of the genera in the Gondwanan clade likely contribute to the instability in the placement of *Columbicola*. Although additional taxon sampling in these genera could be helpful to stabilize the trees, many of these genera are monotypic (e.g. *Dahlemhornia*, *Nesiotinus*, *Trichophilopterus*) such that additional species sampling to break up these long branches will not be possible.

#### Waterbird Lice

The phylogenomic analyses revealed consistent and strong support for a clade of waterbird lice. Although the *Philoceanus*-complex had been identified by prior morphological treatments (Ledger 1980), the close relationships among the other members of this clade had not. Phylogenomic studies of birds (Jarvis et al. 2014, Prum et al. 2015) have revealed a large clade of waterbirds (Aequorlitornithes) that are closely related, including flamingos, grebes, seabirds, cormorants, and herons. These groups all host lice from this waterbird louse clade. Thus, shared ancestry may be involved in the shared ancestry of lice in this group. However, some birds in the waterbird clade (Aequorlitornithes), including penguins, loons, shorebirds, and gulls, do not host lice from this group. In addition, some of the lice in this group occur on waterbirds, such as ducks, coots, and rails, that are not members of this larger avian clade (Aequorlitornithes). Thus, shared habitat by association with water may have facilitated host switching by lice in this group. Furthermore, some raptors, which again are not in this large waterbird clade, also host lice from this group. This may be a case of switching from prey to predators during predation events, which has been shown in the lice of Galapagos Hawks (Whiteman et al. 2004).

Although many of the genera in this clade are wing lice, there are several independent origins of head louse genera in this group. In the case of *Docophoroides*, a head louse from albatrosses, this taxon is sister to a clade (the *Philoceanus*-complex) containing the wing lice of albatrosses. Similarly, *Anatoecus*, a head louse from flamingos and ducks, is sister to the clade containing the wing lice (*Anaticola*) from flamingos and ducks. In other cases, the head lice occur on avian groups that are distantly related to the hosts on which the wing louse sister taxon occurs. For example, the head louse genus *Trabeculus* from tube-nosed seabirds (Procellariiformes) is sister to *Fulicoffula*, a wing louse from coots and rails (Gruiformes).

The genera sampled by more than one species in this clade are all strongly supported as being monophyletic, suggesting generic limits based on morphology in this group may be robust. However, the relationships among genera, particularly the placement of *Ardeicola* is unstable between analyses and some early diverging branches in this clade do not have maximum support. Some genera in this group on long branches, such as *Docophoroides*, *Trabeculus*, *Neophilopterus*, and *Ibidoecus*, are only sampled by a single species. Thus, additional future sampling of other members of these genera may help break up long branches in this and help stabilize relationships among them.

#### Oxilipeurus-*complex*

While members of the *Oxylipeurus*-complex occur mainly on landfowl (Galliformes) throughout the world, one genus, *Splendoroffula*, occurs on turacos in Africa. This group includes both wing lice (*Oxylipeurus* and *Splendoroffula*) and body lice (*Labicotes* and *Chelopistes*). The paraphyly of *Oxylipeurus* (as defined by Price et al. 2003) indicates that further taxonomic examination of this group is needed, likely splitting *Oxylipeurus* into multiple genera. Indeed, such divisions have been recently proposed on the basis of morphology (Gustafsson et al. 2024). While this genus is widespread across most Galliformes, some groups apparently are not host to members of this complex such as New World grouse and Old World quail, partridge, and francolins (Price et al. 2003). Instead, these are prominent hosts of members (*Lagopoecus* and *Cuclotogaster*) of the next group.

#### Otidoecus *and* Degeeriella-*complexes and* Lipeurus

Together the genus *Lipeurus* and the *Otidoecus* and *Degeeriella*-complexes form an extremely well-supported clade with a long stem branch uniting them. The members of these three complexes have long been recognized based on morphological features such as a typically unbroken marginal carina and generalized morphological form. However, we find some unanticipated members of these groups that do contain a broken marginal carina, including *Austrophilopterus* (from toucans) and *Emersoniella* (from kingfishers) that are strongly supported as being within the *Degeeriella*-complex. Generally, the members of these groups occur on land birds, and even the groups of kingfishers that host *Emersoniella* are associated with forested habitats, not water. Both the *Otidoecus* and *Degeeriella* complexes as defined by Ledger (1980) are monophyletic, with *Lipeurus* being sister to the *Otidoecus*-complex. Notably all three of these lineages contain lice hosted by landfowl (Galliformes).

Within the *Otidoecus*-complex, we find a louse from flufftails (Gruiformes: Sarothruridae) closely related to *Cuclotogaster*. There are no prior records of lice from this family of birds, so this might potentially represent a new genus. Known hosts of *Cuclotogaster*, such as francolins (Galliformes), live in the same grassland habitats in Africa as flufftails, so this host-association likely reflects a host-switching event. The other lice in the *Otidoecus*-complex, *Otidoecus* (from bustards) and *Rhynonirmus* (primarily from snipes and woodcocks), also occur on birds in grasslands or wet grasslands, which may again be a case of common habitat association facilitating host-switching among distantly related groups of birds.

We find several cases of paraphyletic genera (as defined by Price et al. 2003) within the *Degeeriella*-complex, and this confirms results of prior studies based on Sanger sequencing data (Johnson et al. 2002, Catanach and Johnson 2015). In some cases, this paraphyly relates to host association. For example, the species of *Degeeriella* from hawks (Accipitridae) are distantly related to those from falcons (Falconidae). Although these two avian groups were previously believed to be closely related, many recent phylogenetic studies have shown that not to be the case (Jarvis et al. 2014, Prum et al. 2015), which can explain why the lice from these two groups are not closely related. In some cases, lice from the *Degeeriella-*complex in different genera, but from the same host group, are closely related, such as *Bucerocophorus*, *Buceroemersonia*, and *Buceronirmus* all from hornbills. In other cases, they are not, such as *Trogoninirmus* from New World trogons and *Trogoniella* from Old World trogons.

#### Struthiolipeurus, Nyctibicola*, Parrot Lice, and the* Brueelia-*complex*

The placement of *Struthiolipeurus* and *Nyctibicola* with a large clade containing parrot lice and the *Brueelia-*complex is somewhat unexpected. Both of these genera had generally unknown affinities based on morphology. In addition, given that paleognathous birds (which includes the ostrich, rhea, and other ratites) comprise the sister taxon to all other birds, the position of *Struthiolipeurus*, which occurs on ostriches and rheas, away from other lice of ratites is noteworthy. This is consistent with previous work that showed that paleognaths likely acquired their lice through host-switching from other birds several times independently (de Moya et al. 2019).

Parrots have been recently shown to be the sister taxon of songbirds (Passeriformes) (Prum et al. 2015), and thus the close relationship of parrot lice with the *Brueelia*-complex (which occurs mainly on songbirds) may reflect the relationships of their hosts. However, the fact that parrot lice form a paraphyletic grade with respect to the *Brueelia*-complex indicates the situation may be more complex than simple codivergence (Johnson and Doña 2024). The earliest divergences within the *Brueelia*-complex occur between lice (*Formicaricola*, *Pseudocophorus*, and *Formicaphagus*) on suboscine passerines and those on mainly oscine passerines plus some other non-passerine birds. Several genera (*Couala*, *Buerelius*, *Meropsiella*, *Meropoecus*, and *Momotmotnirmus*) that occur on non-passerine birds form a clade sister to those that occur primarily, but not exclusively, on oscine passerines. Members of this complex are morphologically diverse, including wing lice, head lice, and body lice ecomorphs, as well as more generalized forms (Bush et al. 2016). Previous studies based on Sanger sequence data sets indicated that congruence between host and parasite phylogenies was variable within this group and related to these ecomorphological differences (Sweet et al. 2018). However, many nodes along the backbone of these trees were weakly supported given the small number of genes sampled (Bush et al. 2016). Given the high species diversity of this group (containing well over 10% of all species of feather lice), and the large number of genera recently described (Gustafsson and Bush 2017), additional sampling within the *Brueelia*-complex is needed to fully uncover the patterns of diversification in this group.

#### Penenirmus, Quadraceps*, and* Philopterus-*complexes*

Together with some other related genera, the *Penenirmus*, *Quadraceps*, and *Philopterus*-complexes form a major highly supported clade in the tree. Although clearly members of this clade, the exact position of some enigmatic genera (*Vernoniella*, *Furnariphilus*, and *Colilipeurus*) within this clade is somewhat unstable. These genera occur on distantly related groups of birds (anis, antbirds, and mousebirds respectively) from different geographic regions (South America and Africa), so it is currently unclear what factors may have been involved in the diversification of these lineages. While in this tree, *Turnicola* falls within the genus *Penenirmus* (as defined by Price et al. 2003), broader sampling of these two genera in a prior phylogenomic study (Johnson et al. 2021) revealed that the genus *Penenirmus* (*sensu lato*) is likely to be monophyletic. Recent work has split genera from within *Penenirmus*, recognizing *Laimoloima* (Gustafsson et al. 2022) from Old World barbets and *Picophilopterus* primarily on woodpeckers and allies (Piciformes). However, there are additional phylogenetic complexities (Johnson et al. 2021) that may require description of additional genera under this framework.

We find strong support for a clade containing the *Quadraceps*-complex (*Cummingsiella*-complex of Ledger 1980), but also including additional genera (*Ardeiphagus*, *Lunaceps*, *Incidifrons*, and *Saemundssonia*). Most of the members of the *Quadraceps*-complex occur on the avian order Charadriiformes (shorebirds, gulls, and allies), but there are several cases of members on different groups of birds such as herons, kingfishers, rails, hamerkop, and cranes. All members of this group tend to be associated with water, and this shared habitat association may have again facilitated host-switching.

Within the *Quadraceps*-complex there is a deep divergence between *Cirrophthirius* (from avocets) and all other members of the group, which form a strongly supported clade. Although most members of this group are wing lice or more generalized forms, head lice have evolved at least two times independently (*Incidifrons* and *Saemundssonia*). Both of these genera appear to be monophyletic, even though *Saemundssonia* occurs on two different orders of birds. The genus *Quadraceps* (as defined by Price et al. 2003) is highly paraphyletic, with most of the other genera in this complex being in multiple derived positions within the *Quadraceps* genus. It is likely that extensive taxonomic revision of this complex is needed to make generic boundaries reflect evolutionary relationships.

All genera in the *Philopterus*-complex, broadly defined, are of the head louse ecomorph. There are three main lineages within this group, *Alcedoecus* (from kingfishers), *Philopterus* (mainly *sensu* Price et al. 2003), and a clade containing *Strigiphilus*, *Craspedorrhynchus*, *Cuculoecus*, and *Debeauxoecus* (formerly placed in *Philopterus*, e.g. by Price et al. 2003 but subsequently recognized to be a distinct lineage using Sanger sequencing data [Kolencik et al. 2022]). Within this latter clade, *Strigiphilus*, *Craspedorrhynchus*, and *Cuculoecus* are all from unrelated groups of birds of prey (owl, hawks, and falcons) or brood parasites (cuckoos), and this ecology is expected to be a factor that would facilitate host switching. However, the only other genus in this clade (*Debeauxoecus*) is from pittas, a comparatively small group of suboscine passerines that only occur in the Old World. Thus, if ecologically facilitated host-switching was involved in the diversification of this clade, it may have involved lineages of hosts that are currently extinct.

Within the genus *Philopterus* (as defined by Price at al. 2003), there are several distinct lineages, some of which are recognized as distinct genera by some authors. Notably, the lineage hosted by jacamars and puffbirds has been placed in *Mayriphilopterus* (Mey 2004) and this is the earliest diverging lineage in this group. Two other lineages hosted by non-passerines (*Clayiella* on motmots and *Vinceopterus* on trogons) also together form a group and this is sister to a clade containing the lice from suboscine passerines, which some recognize as the genus *Tyranniphilopterus*. Although somewhat limited by taxon sampling, we find a clade containing all the members of *Philopterus* occurring on oscine passerines, with a species occurring on wood-hoopoes embedded within it. This clade has also been split by some authors (Mey 2004) into several genera (including *Philopteroides*, *Australophilopterus*, *Corcorides*, *Paraphilopterus*), and indeed we find support for monophyly of a clade containing members of *Philopterus sensu stricto*. However, our results together with results of a Sanger sequencing data set (Kolencik et al. 2022) suggest generic reclassification within this clade might be premature until more taxon sampling can be conducted using genome scale data sets and more detailed morphological investigation.

#### Goniodidae

Sometimes classified as the *Goniodes*-complex (Ledger 1980), this group of feather lice has often been considered as one deserving recognition at the family level (Smith 2000, Johnson et al. 2001, 2011). Members of this clade occur on landfowl (Galliformes) and pigeons and doves (Columbiformes) and are all of the body louse ecomorph. Previous work based on Sanger sequencing data sets (Johnson et al. 2001) have provided evidence that the genus *Chelopistes*, which has strong morphological affinities with members of Goniodidae, is distantly related. Our current study strongly supports this conclusion, with the genera *Chelopistes* and *Labicotes* being more closely related to *Oxylipeurus* and not to members of Goniodidae. A Sanger dataset with more extensive taxon sampling (Johnson et al. 2011) indicated that members of this group from doves are embedded within those from landfowl, implying switching from landfowl to doves. However, this study also indicated a potential case of switching from doves back to landfowl. The current phylogenomic study lacks the taxon sampling to test this hypothesis, but we do find strong support for lice from doves being a monophyletic clade embedded within those from landfowl as in the prior study. A recent phylogenomic study with increased taxon sampling also finds evidence of the switch from landfowl to doves, but not for a switch back to landfowl (Sweet et al. 2025). We do not find support for monophyly of either *Goniodes* or *Goniocotes* (sensu Price et al. 2003), and some authors have erected additional genera (Mey 2009) for some lineages. Multiple additional genera have also been suggested for lineages of lice within Columbiformes (e.g. Tendeiro 1969), but the current study does not have enough taxon sampling of these lineages to form any taxonomic conclusions regarding these genera. Additional taxon sampling of lice within this group from both host groups is needed (along the lines of Sweet et al. 2025), in combination with morphology, to help provide a reclassification of the genera in this group.

#### Caracaricola, Podargoecus, and Mulcticola

A strongly supported branch unites the genera *Caracaricola*, *Podargoecus*, and *Mulcticola*. These genera are rather disparate in gross morphology, with *Caracaricola* having a form like a generalist ecomorph, *Podargoecus* much like a head louse ecomorph, and *Mulcticola* the form of a wing louse ecomorph. *Podargoecus* is hosted by frogmouths and *Mulcticola* by nightjars, two lineages within the same broader group of birds (Caprimulgiformes, in part). However, *Caracaricola* is hosted by caracaras (a group within Falconiformes) which are very distantly related. Unlike falcons, which are predators, caracaras are scavengers. This scavenging behavior might have facilitated host switching between an ancestral nightjar or relative and an ancestor of caracaras. Future studies should investigate if there are any morphological characters that might unite these three genera, given they form such a strongly diagnosed clade in the phylogenomic trees.

#### South American clade plus Rallicola

A number of genera exclusive to South America and the genus *Rallicola* appear together in a strongly supported major clade. The exclusively Neotropical members of this clade include three genera of tinamou lice (*Tinamotaecola*, *Pseudolipeurus*, and *Pseudophilopterus*), two genera of hoatzin lice (*Pessoaella* and *Osculotes*), and two newly described genera of trumpeter lice (*Palmaellus* and *Psophiicola*). The genus *Rallicola* occurs on six orders of birds throughout the world. However, this genus has also undergone extensive diversification on Neotropical hosts, including woodceepers and some Neotropical woodpeckers. The genus *Rallicola* also occurs on kiwis (New Zealand), coots and rails (worldwide), coucals (Old World tropics), jacanas (worldwide tropics), and the Marianas Island crow. Notwithstanding this incredibly broad biogeographic and host distribution, the monophyly of the genus *Rallicola* (as defined by Price et al. 2003) is strongly supported, being united on a long branch. The genus *Psophiicola* was synonymized into *Rallicola* by Price et al. (2003). Although this would still leave *Rallicola* monophyletic, *Psophiicola* is on a long branch sister to and well separated from the remaining members of *Rallicola* in our tree, further justifying its status as a unique genus.

In this broader clade, there are multiple examples of lice of different ecomorphs, but from the same group of hosts being each other’s closest relatives. Within the three genera of lice from tinamous, the genus *Pseudophilopterus* (a head louse) is embedded within *Pseudolipeurus* (a wing louse). Together these two genera are closely related to *Tinamotaecola*, a generalist louse from tinamous, and this was also previously shown by Virrueta Herrera et al. (2020). Further sampling of these genera will be needed to unravel the unexpected result of paraphyly of *Pseudolipeurus* with respect to *Pseudophilopterus* and whether there are morphological characters that may also reflect these relationships. Similarly, results on the lice of hoatzins also indicate that a more generalized louse, *Pessoaella*, is sister to a morphologically highly disparate body louse, *Osculotes*. Although *Psophiicola*, a generalized louse, is not sister to the head louse from trumpeters (*Palmaellus*), together they form a grade with respect to *Rallicola*. Thus, in general this broader clade of lice appears to be highly adaptable, both in terms of moving into new niches on the same host but also diversifying across distantly related groups of birds as is the case in the genus *Rallicola*.

#### Heptapsogastridae

One of the most distinctive groups of avian feather lice is the group Heptapsogastridae, which is primarily found on tinamous in the Neotropics. This family has been separated from Philopteridae by many (Carriker 1936, Hopkins and Clay 1952), but not all (Ward 1957, Price et al. 2003), workers based on unique, although sometimes misinterpreted, characters of the abdomen (reviewed by Smith 2000). Monophyly of this family is strongly supported, in agreement with a prior phylogenomic study (Virrueta Herrera et al. 2020) and morphological work (Smith 2000). Two lice from seriemas, an exclusively South American group of birds, are deeply embedded within those from tinamous, being in the same genus *Heptapsogaster*, as other lice of tinamous. Presumably this represents a host-switch from tinamous to seriemas. Although we did not have multiple sampling of many genera of Heptapsogastridae, we did find that *Lamprocorpus* was embedded within *Heptapsogaster* (*sensu* Price et al. 2003). Further work with expanded taxon sampling is needed to explore these relationships in more detail.

### Patterns of Diversification

Although usually based in morphological diagnoses, prior taxonomic practices in feather lice have often circumscribed genera of lice based on major host groups, such as order or family of birds. Broad ecomorphological differences also appear to be key determinants of generic boundaries. These taxonomic decisions notwithstanding, there are additional broad patterns of host association across the tree of feather lice that indicate biological underpinnings of evolutionary diversification. Consistent with prior studies (de Moya et al. 2019), the lice of paleognath birds (ratites and tinamous) are spread throughout the tree, in several cases being in highly derived positions, such as the lice of kiwis and tinamous. Paleognaths are the sister taxon of all other birds, diverging from them before the Cretaceous-Paleogene (K-Pg) boundary (Jarvis et al. 2014, Prum et al. 2015). Thus, it appears the modern diversification of feather lice did not originate on the common ancestor of all birds (de Moya et al. 2019).

The next group of birds to diverge, again before the K-Pg boundary (Prum et al. 2015), is the Galloanserae, a group containing waterfowl (Anseriformes) and landfowl (Galliformes). This group is the sister taxon to all other neognathous birds (i.e. Neoaves), and Neoaves underwent extensive radiation right after the K-Pg boundary (Prum et al. 2015). Lice from Galloanserae occur in several of the earliest diverging major clades of lice. In particular, the Gondwanan clade contains lice from early diverging lineages of waterfowl, such as screamers and magpie geese. Lice from waterfowl also occur in the waterbird louse clade near the base of the tree, but this may have been switching from flamingos to ducks (Johnson et al. 2006), rather than reflecting the early divergence of waterfowl. The next two clades, the *Oxylipeurus*-complex and *Lipeurus* plus the *Otidoecus* and *Degeeriella*-complexes prominently feature lice of landfowl. Prior cophylogenomic analyses on a smaller taxon set (de Moya et al. 2019) indicated that feather lice may have begun their diversification on the common ancestor of waterfowl and landfowl, and our current tree with expanded taxon sampling appears to be consistent with this conclusion.

We also find several cases in which different genera of lice from the same host group are closely related. This pattern occurs in all the major clades outlined above. In several cases, this involves lice of different ecomorphs from the same host group being closely related, a pattern initially outlined by Johnson et al. (2012). With the expanded taxon sampling in this study, such situations do not always involve terminal sister pairs of genera, such as *Anatoecus* (head louse) and *Anaticola* (wing louse) from flamingos and ducks; *Ibidoecus* (head louse) and *Ardeicola* (wing louse) from ibises; and *Palmaellus* (head louse) and *Psophiicola* (generalist louse) from trumpeters. In these cases, such genera are still extremely close relatives being separated by only one or two branch points. There still remain many cases in which genera of lice of different ecomorphs from the same host lineage are indeed terminal sister taxa, such as *Echinophilopterus* (head louse) and *Psittoecus* (body louse) from parrots; *Paragoniocotes* (body louse) and *Neopsittaconirmus* (wing louse) from parrots; *Pseudophilopterus* (head louse) and *Pseudolipeurus* (wing louse) from tinamous; and *Pessoaella* (generalist louse) and *Osculotes* (body louse) from hoatzin. Thus, the general pattern of ecomorphological divergence of lice within host lineage, but convergence across lineages (Johnson et al. 2012), appears to be borne out, albeit with more complexity than previously indicated.

There are also some cases of generic diversification of related groups of lice within a host lineage, but without ecomorphological diversification related to how lice escape preening (i.e. head louse, wing louse, body louse). One example is the *Philoceanus*-complex, which has radiated into almost a dozen genera across tube-nosed seabirds, particularly albatrosses. These genera are of the wing louse body form. The long wings of albatrosses may have allowed lice to radiate into different specific niches across the wing. Another prominent example is the *Goniodes*-complex, which consists entirely of lice of the body louse ecomorph. However, within this complex, both the lice of landfowl and of doves are divided into multiple genera. This division is primarily related to the body size of these lice, and in fact it is not clear if the genera *Goniodes* and *Goniocotes* are separable by anything other than body size. Thus, these body lice may be partitioning niches on the body of the bird, possible relating to whether they tend to reside on the dorsal or ventral side of the feather (Johnson et al. 2005). Perhaps the most prominent example of such a radiation is the body lice of tinamous, Heptapsogastridae (Virrueta Herrera et al. 2020). These lice have radiated into 15 genera, up to 8 of which can sometimes coexist on the same host species (Price et al. 2003). It is still unclear what factors may have facilitated this extensive *in situ* radiation on tinamous, but may relate to the fact that they do not have downy feathers in their apteria (the space between feather tracts) (Chandler 1916, Bailey 1955), unlike other birds which do.

In addition to radiation of louse lineages within a host group, there are also cases of lice of different ecomorphs from the same host group being very distantly related. For example, the wing lice (*Columbicola*) and body lice (Goniodidae) of doves are separated in two very distantly related lineages in the tree. The wing lice of doves radiated after their hosts (Boyd et al. 2022) and it appears their body lice are derived from within the lice of landfowl, possibly representing a recent host switch (Sweet et al. 2025). Thus, doves may have been an open niche for feather lice that might have facilitated radiation across these hosts. Similarly, the wing lice (*Fulicoffula*), head lice (*Incidifrons*), and generalists (*Rallicola*) of coots and rails are in three very distinct, distantly related clades. Like doves, coots and rails have very high dispersal capabilities being found on many isolated Pacific Islands. As has been shown for the lice of doves (Boyd et al. 2022), it may be that high dispersal ability places lineages of birds into novel patterns sympatry, which may in turn facilitate switching of lice between host lineages. Finally, the lice of songbirds (Passeriformes) are scattered throughout many clades in the tree, including the *Degeeriella*-complex, *Brueelia*-complex, *Philopterus*-complex, *Penenirmus*-complex, and *Rallicola*. This is perhaps not surprising given the fact that songbirds form the majority (around two-thirds) of avian diversification. Generally, songbird lice of these complexes are found on some, but not all, lineages of songbirds, indicating that interspecific competition between lice on songbirds might restrict the host distribution of some of these groups of lice.

## Acknowledgements

We thank C. Adam, R.J. Adams, A. Alexio, S. Barker, J. Bates, B. Benz, S. Bertelli, I. Beveridge, S. Bush, S. Cameron, H. Campbell, M. Carbajal, T. Catanach, J. Cheriton, T. Chesser, D. Clayton, M. Cottam, L. Cueto, F. Daunt, B. Dawson, E. Diblasi, H. Eascott, R. Empson, H. Fandel, R. Faucett, R. Furness, A. Garitano-Zavala, T. Galloway, N. Gawani, T. Gnoske, S. Goodman, A. Gouvea, D. Gustafsson, J. Hagelin, C. Harbison, N. Hoffman, R. Jakob-Hoff, J. Jankowski, J. Jolly, R. Junge, J. Kaderitz, S. Kenney, J. Kirchman, J. Klicka, A. Krater, E. Kuschel, D. Lane, P. Loi, H. Lutz, B. Marks, J. Malenke, I. Mason, K. McCracken, J. Merkel, M. Meyer, M. Miller, R. Moyle, E. Neutze, B. O’Shea, E. Osnas, R. Palma, R. Palmer, V. Piacentini, A. Porzecanski, D. Roach, M. Robbins, K. Rose, J. Sailer, D. Santiago-Alarcon, J. Scherer, F. Schunck, F. Sheldon, V. Smith, S. Sontshugen, D. Steadman, A. Sweet, S. Trewick, W. Veronesi, D. Verrier, S. Volponi, J. Weckstein, N. Whiteman, D. Willard, R. Wilson, B. Winger, C. Witt, J. Wombey, B. Zonfrillo for assistance in obtaining specimens for this study. We thank A. Hernandez and C. Wright at the University of Illinois Roy J. Carver Biotechnology Center for assistance with Illumina sequencing. This work was supported by U.S. NSF DEB-1239788, DEB-13426045, DEB-1926919, DEB-1925487, and DEB-2328118 to K.P.J.

## Supplementary files

**Table S1**. List of samples included in this study, with associated host species, locality information, sequencing, and assembly details.

**Figures S1.** ASTRAL phylogenetic tree of ischnoceran feather lice and relatives. Branch support values represent local posterior probabilities

**Figure S2.** Biogeographic states recovered in the BioGeoBEARS ancestral area reconstruction. Simple areas represent the five major regions (NW, AN, EA, AF, AU), while multi-area states denote combinations of these regions.

## Notes

https://doi.org/10.6084/m9.figshare.29266997

### Competing Interest Statement

The authors have declared no competing interest.

