## Supplementary material for "A phylogenomic framework for avian feather lice (Phthiraptera: Ischnocera)": Figure S1

Figure S1. ASTRAL phylogenetic tree of Ischnoceran feather lice and relatives. Branch support values represent local posterior probabilities

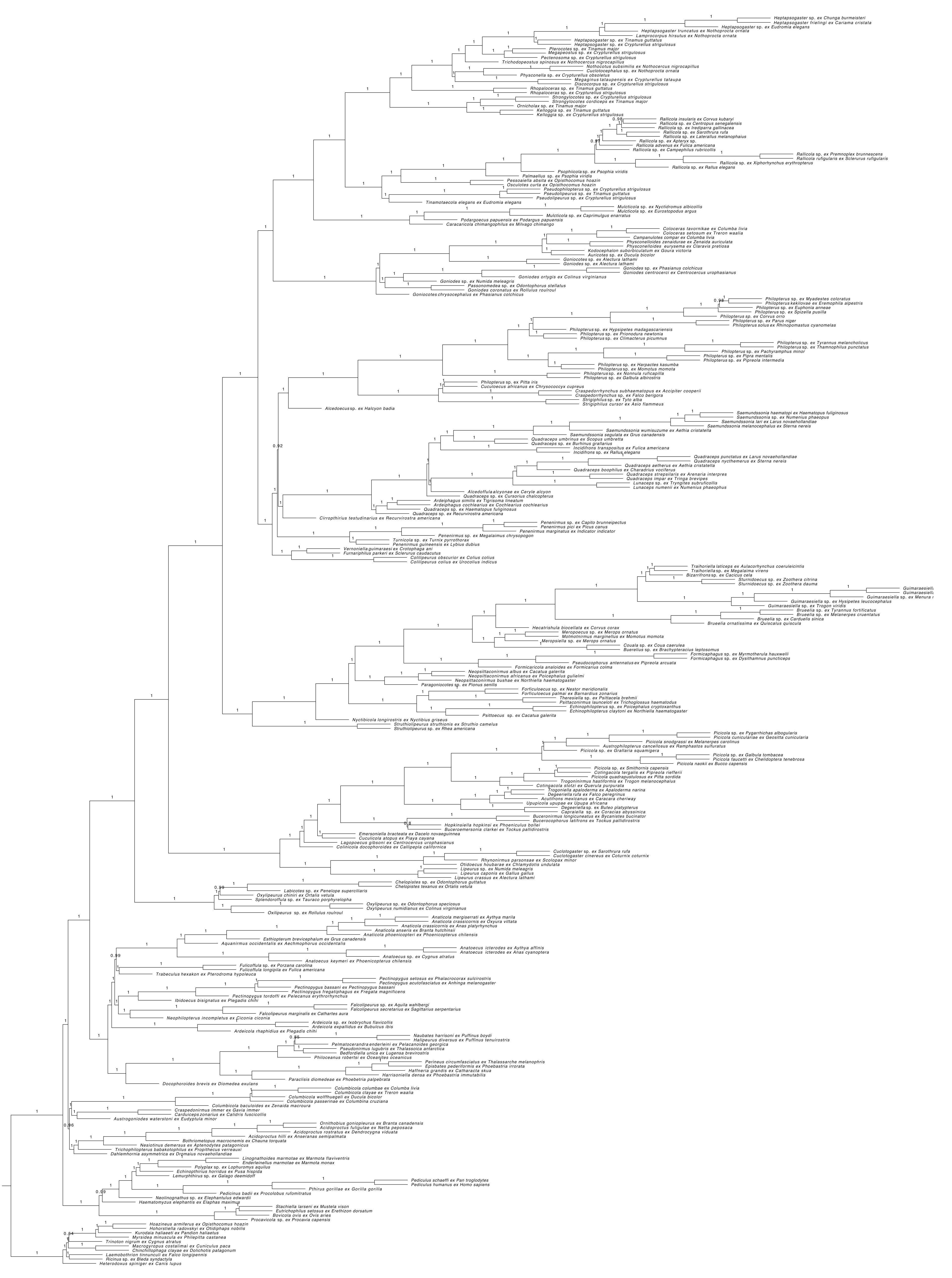
