## Supplementary material for "A phylogenomic framework for avian feather lice (Phthiraptera: Ischnocera)": Figure S2

Figure S2. Biogeographic states recovered in the BioGeoBEARS ancestral area reconstruction.
Simple areas represent the five major regions (NW, AN, EA, AF, AU), while multi-area states denote combinations of these regions.

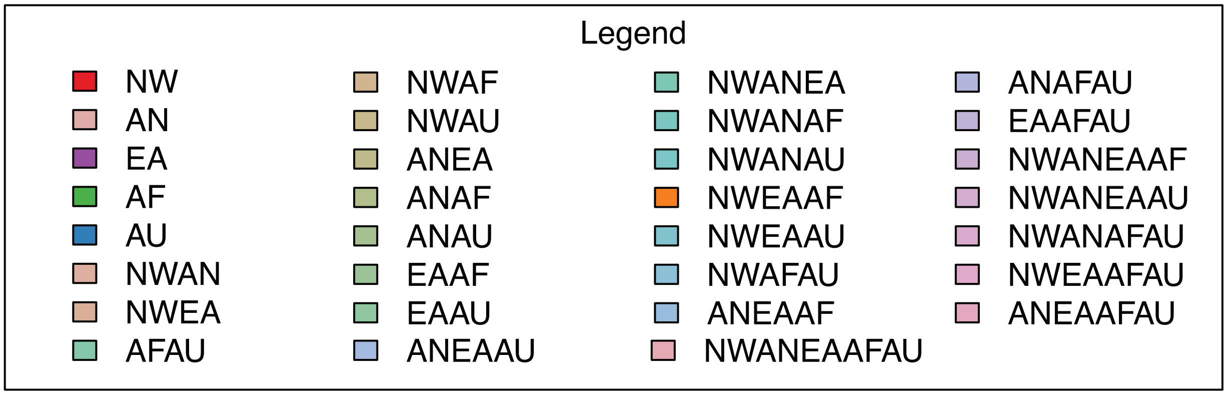
